# Evolution of thermal physiology alters the projected range of threespine stickleback under climate change

**DOI:** 10.1101/2021.02.25.432865

**Authors:** Sara J.S. Wuitchik, Stephanie Mogensen, Tegan N. Barry, Antoine Paccard, Heather A. Jamniczky, Rowan D.H. Barrett, Sean M. Rogers

**Affiliations:** Department of Biological Sciences, University of Calgary, 2500 University Dr NW, Calgary, AB, T2N 1N4, CANADA; Informatics Group, Harvard University, 52 Oxford St, Cambridge, MA, 02138, USA; Department of Biology, Boston University, 5 Cummington Mall, Boston, MA, 02215, USA; Redpath Museum and Department of Biology, McGill University, 845 Sherbrooke St W, Montreal, QC, H3A 0G4, CANADA; Department of Cell Biology & Anatomy, Cumming School of Medicine, University of Calgary, 3330 Hospital Dr NW, Calgary, T2N 4N1, CANADA; Bamfield Marine Sciences Centre, 100 Pachena Rd, Bamfield, BC, V0R 1B0, CANADA

**Keywords:** thermal biology, critical thermal limits, evolution, species distribution models, climate change, threespine stickleback

## Abstract

Species distribution models (SDMs) are widely used to predict range shifts but could be unreliable under climate change scenarios because they do not account for evolution. The thermal physiology of a species is a key determinant of range and thus incorporating thermal trait evolution into SDMs might be expected to alter projected ranges. We identified a genetic basis for physiological and behavioural traits that evolve in response to temperature change in natural populations of threespine stickleback *(Gasterosteus aculeatus).* Using these data, we created geographic range projections using a mechanistic niche area approach under two climate change scenarios. Under both scenarios, trait data was either static (‘no evolution’ models), allowed to evolve at observed evolutionary rates (‘evolution’ models), or allowed to evolve at a rate of evolution scaled by the trait variance that is explained by quantitative trait loci (QTL; ‘scaled evolution’ models). We show that incorporating these traits and their evolution substantially altered the projected ranges for a widespread panmictic marine population, with over 7-fold increases in area under climate change projections when traits are allowed to evolve. Evolution-informed SDMs should improve the precision of forecasting range dynamics under climate change, and aid in their application to management and the protection of biodiversity.

## Introduction

Temperature is a powerful driver of global biogeography and species distributions frequently reflect temperature gradients in both aquatic and terrestrial habitats (Hochachka & Somero, 2002). Many species adopt thermal strategies (such as thermoregulation or acclimation) that determine their thermal niche (Coutant, 1987; Huey & Kingsolver, 1989; Huey & Slatkin, 1976) and thermal traits can provide a target for directional selection if the environment changes to include temperatures outside the range encompassed by the thermal niche. Adaptation can thus permit species to persist at temperatures that would have previously led to extirpation (Hoffman & Sgrò, 2011; Sexton et al., 2009). Under moderate climate change scenarios, mean global oceanic temperature is projected to increase in excess of 2 °C by the end of the century (IPCC, 2014), with more extreme changes predicted in localized regions (Eyer et al., 2019; Walther et al., 2002) for both cooling and warming events (Hu et al., 2018). Accurately predicting species range patterns under climate change therefore requires data for temperature-associated adaptive trait evolution (Bay et al., 2017; Bush et al., 2016; Catullo et al., 2015).

The genetic basis of thermal traits has been explored in many terrestrial ectotherms (Rolandi et al., 2018; Angilletta et al., 2002; Bowler & Terblanche, 2008; Dillon et al., 2009) and there have been integrative frameworks proposed for how best to use this information to assess the vulnerability of organisms to environmental changes (Araújo et al., 2013; Bay et al., 2017; Huey et al., 2012). Given the differences in thermal biology between terrestrial and aquatic ectotherms (Sunday et al., 2011) and differential warming between terrestrial and aquatic habitats (IPCC, 2014, 2018), predicting how the thermal biology of aquatic organisms will respond to environmental changes requires an understanding of the genetic architecture underlying ecologically relevant thermal traits (Healy et al., 2018) in aquatic systems. The genetic basis of thermal traits has been explored in a number of socioeconomically and culturally important fish species (Everett & Seeb, 2014; Jackson et al., 1998; Jin et al., 2017; Larson et al., 2016; Muñoz et al., 2014; Perry et al., 2001) and there has been an extensive body of work on the thermal biology of the common killifish *(Fundulus heteroclitus,* Bryant et al., 2018; Fangue et al., 2006, 2009; Healy et al., 2018). These studies suggest that a significant proportion of phenotypic variance in thermal traits can be explained by genetic variation *(e.g.,* 35 SNPs explained 51.9% of the variation for upper thermal tolerance in killifish, Healy et al., 2018), highlighting the potential for heritable responses to changes in thermal conditions.

While the climate and ocean temperatures are warming overall, thermal events characterized by both cold and heat extremes are occurring with increasing frequency (Hu et al., 2018; IPCC, 2014, 2018; Stott, 2016). Regionally downscaled environmental changes can be highly spatially heterogenous relative to global temperature trends, and are likely to be the most relevant for species-specific responses in the context of contemporary climate change (Walther et al., 2002). For example, range expansions of marine species have been mediated by both ocean-scale warming and regional cooling (Zeidberg & Robison, 2007). In such cases, population persistence and adaptation will rely on both upper and lower critical thermal limits, forcing organisms to respond independently to extreme heat and extreme cold in separate events (Herring et al., 2015, 2018, 2020; Hu et al., 2018; Walther et al., 2002). Importantly, adaptation in one thermal trait can shift the thresholds of correlated thermal traits (Buckley & Huey, 2016; Denny & Dowd, 2012; Hoffman & Sgrò, 2011; Huey & Kingsolver, 2011). Associations between traits influence the rate of trait adaptation and thus correlated traits must be considered when attempting to predict future species distributions under climate change (Bestion et al., 2015; Hoffman & Sgrò, 2011; Valladares et al., 2014). Recently there have been steps to incorporate theoretical trait evolution into SDMs *(e.g.,* by including the breeder’s equation for a key phenotypic trait, Bush et al., 2016; Catullo et al., 2015; Kearney & Porter, 2009). However, no model has used empirical estimates of evolutionary rates to inform projected species ranges under climate change scenarios.

Threespine stickleback *(Gasterosteus aculeatus,* Fig. 1a) are a useful vertebrate species for understanding the impact of adaptation on range dynamics under climate change. This species exhibits widespread phenotypic variation (Hendry et al., 2013), there are publicly available genomic resources (Jones, Grabherr, et al., 2012), and there is prior research on temperature-associated evolutionary rates for this species (Barrett et al., 2011; Morris et al., 2018). It is also feasible to artificially breed multiple hybrid generations in common garden lab environments. Moreover, there is a regionally downscaled climate model for the Pacific Northwest (Alexander et al., 2018; IPCC, 2014; Reynolds et al., 2007), which makes panmictic marine populations of *G. aculeatus* in the eastern Pacific Ocean (Kirch et al., 2021; Morris et al., 2018) an ideal system for testing how adaptive trait variation could affect projections of species range under climate change.

**Figure 1.**
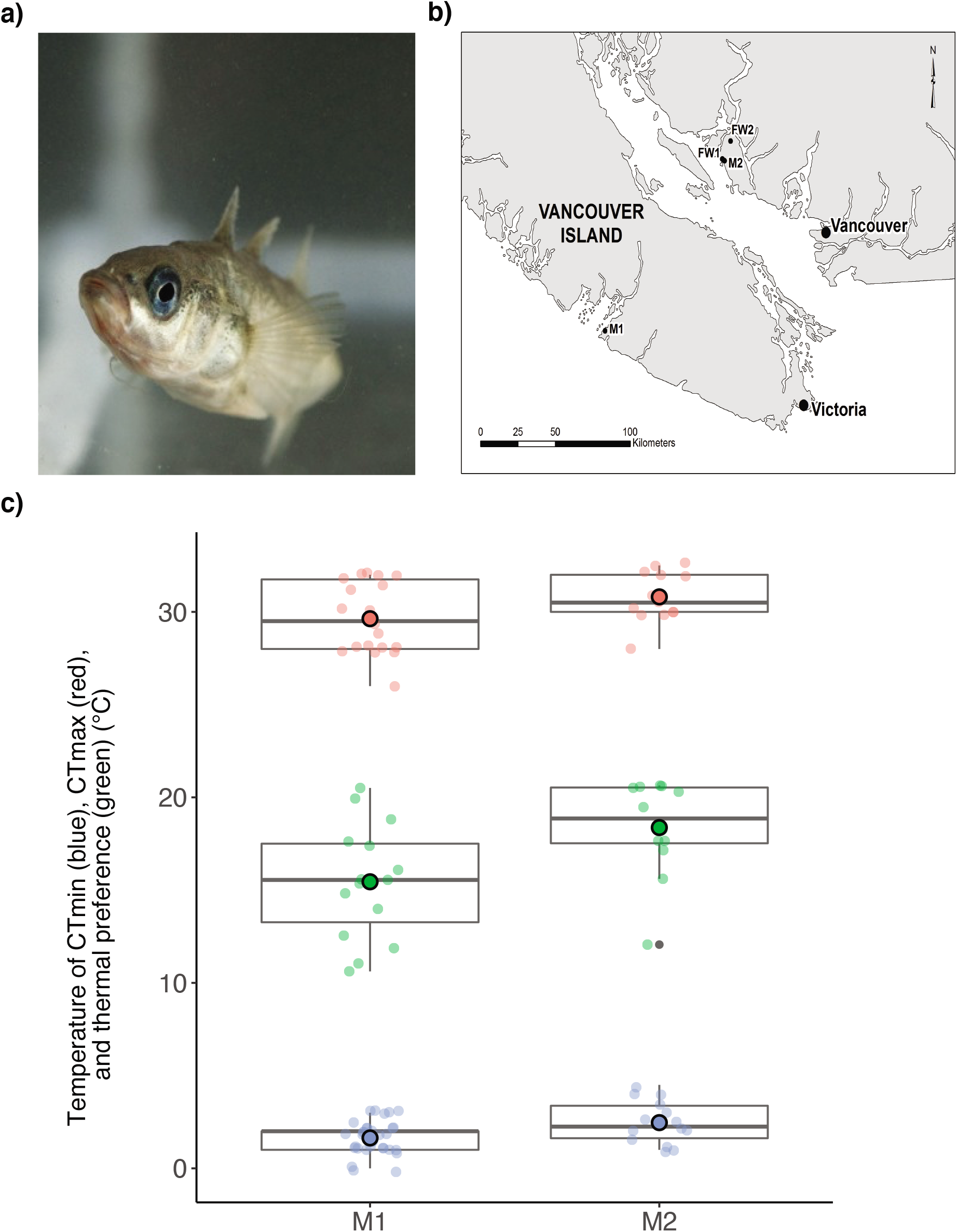
a) Adult threespine stickleback *(Gasterosteus aculeatus)* from a single genetic cluster were sampled from b) two marine and two freshwater populations in the Canadian Pacific Northwest. These populations were assayed for c) thermal preference (green), critical thermal minima (CTmin, blue) and maxima (CTmax, red). Thermal trait values for marine populations (M1 and M2) were incorporated into the species distribution models.

The objective of this study is to incorporate ecologically and evolutionarily relevant data on thermal traits into end-of-century projections of species distribution models. To accomplish this, we assessed physiological and behavioural temperature-associated traits in two marine and two freshwater populations of threespine stickleback, as well as within-population F1 families and marine-freshwater hybrid F1 and F2 families. We used the hybrid families to construct genetic linkage maps and identified multiple quantitative trait loci associated with thermal preference, as well as upper and lower critical thermal limits. Using these genetically-based traits and empirically derived evolutionary rate data for this species (Barrett et al., 2011), as well as others (Morgan et al., 2020; Sanderson et al., 2021), we constructed species distribution models using a mechanistic niche area approach that integrated the trait data reported in this study and allowed those traits to evolve under two climate change scenarios, three distinct types of evolutionary models, and three evolutionary rates.

## Materials and Methods

### Sample collection and husbandry

We collected adult *Gasterosteus aculeatus* from two marine populations (Bamfield, M1, 48°49’12.69”N 125° 8’57.90”W; Garden Bay Lagoon, M2, 49°37’52.84”N 124° 1’49.26”W) and two freshwater populations (Hotel Lake, FW1, 49°38’26.94”N 124° 3’0.69”W; Klein Lake, FW2, 49°43’32.47”N 123°58’7.83”W) in southwestern British Columbia (Fig. 1b) at less than 0.5 m depths. To ensure that any genetic variation identified would be relevant for adaptation within this region, we confined our study to a single genetically panmictic cluster of populations (Morris et al., 2018). Individuals were maintained in a flow-through system and under a photoperiod that mimicked the natural source population environment during collection periods before transport. We transported the fish to the Life and Environmental Sciences Animal Resources Centre at the University of Calgary, where we separated the fish into population-specific 113 L glass aquaria at a density of approximately 20 fish per aquarium. We acclimated marine individuals to freshwater salinity over one week and maintained fish in a common environment (salinity of 4-6 ppt, water temperature of 15 ± 2 °C, and a photoperiod of 16L:8D). Individuals were allowed to acclimate for at least 2 weeks before experiments (1 week for stress reduction post-transfer, 1 week for common garden environment acclimation and salinity ramp). Each common garden aquarium was on a closed system with individual filters, air stones, and water supply. We fed all adult fish *ad libitum* once per day with thawed bloodworms (Hikari Bio-Pure Frozen Bloodworms). All collections and transfers were approved by the Department of Fisheries and Oceans (marine collections and provincial transfers), the Ministry of Forests, Lands, and Natural Resource Operations (freshwater collections), and the Huu-ay-aht First Nations (marine collections).

### Crossing design for marine and freshwater F1 families

We collected eggs from females and fertilized the eggs with extracted testes from euthanized males. We transferred the fertilized egg mass to a mesh-bottomed egg incubator suspended in a 37 L aquarium for hatching. Each hatching aquarium was maintained with a single air stone and a filter. Once hatched, we reared the larval fish in 37 L hatching aquaria until they reached a total length (TL) of approximately 1 cm, after which we split the families into family-specific 113 L aquaria to maintain suitable densities. We fed the larval fish *ad libitum* twice daily with live *Artemia spp.* nauplii, and then gradually transitioned the diet to chopped, thawed bloodworms (Hikari Bio-Pure Frozen Bloodworms) *ad libitum* once daily as they reached approximately 2 cm TL. The F1 families were maintained in a common garden environment identical to that of the F0 populations. We produced one F1 family for each population (M1_F1, M2_F1, FW1_F1, and FW2_F1).

### Crossing design for hybrid mapping families

To generate genetically heterogeneous marine-freshwater F1 families from wild F0 parents, we collected eggs from marine females and fertilized the eggs with extracted testes from euthanized freshwater males. Egg masses were hatched, and juveniles were reared, as detailed above. We produced one F1 family of M1xFW1 hybrids (hereafter referred to as H1_F1) and three F1 families of M1xFW2 hybrids (hereafter referred to as H2_F1). The hybrid F1 families were maintained in a common garden environment identical to that of the F0 populations. To generate F2 families for linkage map construction and quantitative trait loci (QTL) mapping, we crossed siblings from the same F1 family using the same methodology that was used to generate the F1 families. The amount of trait variation explained by these hybrid crosses is likely to be higher than would be observed in marine-marine or freshwater-freshwater crosses, increasing our likelihood of detecting QTL. Overall, we produced one F2 family of H1xH1 hybrids (referred to as H1_F2) and three families of H2xH2 hybrids (referred to as H2_F2_1, H2_F2_2, and H2_F2_3). All F2 individuals were raised as described above in a common garden environment identical to that of the F0 and F1 individuals to ensure consistent thermal history.

### Thermal tolerance and preference experiments

To assess the lower and upper limits of physiological thermal tolerance, we conducted standard critical thermal minimum (CTmin) and maximum (CTmax) experiments on adult fish (Barrett et al., 2011; Fangue et al., 2006; Hutchison, 1961). At these sublethal limits, the fish experience a loss of equilibrium (LOE) at which they lose the ability to escape conditions that would ultimately lead to their death (Beitinger et al., 2000). Before each experiment, individuals were fasted for 24 hours. Our experimental tank held 1000 mL glass beakers aerated individually to prevent thermal stratification. After a 15-minute acclimation to the experimental apparatus in the individual beakers, we cooled or heated the water (for CTmin or CTmax, respectively) at a rate of approximately 0.33 °Cmin^-1^. We assessed wild F0 individuals (*n_M1_* = 32, *n_M2_* = 14, *n_FW1_* = 15, *n_FW2_* = 16, *N* = 77; Fig. S2), lab raised F1 (*n_M1_F1_* = 13, *n_M2_F1_* = 15, *n_FW1_F1_* = 15, *n_FW2_F1_* = 15, *N* = 58; Fig. S2), and F2 individuals (*n_H1_F2_* = 28, *n_H2_F2_1_* = 36, *n_H2_F2_2_* = 21, *n_H2_F2_3_* = 17, *N* = 102; Fig. S3). All individuals were assessed for CTmin, allowed to recover for at least three days, then assessed for CTmax to keep thermal stress history consistent. The onset of erratic behaviours associated with a behavioural stress response (i.e., ‘agitation windows’, Turko et al., 2020) occurred below 5.0 °C and above 25.0 °C during CTmin and CTmax experiments, respectively. Normal behaviour was observed between 5.0 °C and 25.0 °C, whereas outside of those temperatures, individuals gradually exhibited more extreme stress responses (*e.g.,* increased gilling rate, erratic movement, muscle spasms, listing, as outlined by the Canadian Council of Animal Care guidelines) until reaching LOE and the inability of an individual to right itself (the experimental endpoint, measured in 0.5 °C increments, Barrett et al., 2011; Fangue et al., 2006; Hutchison, 1961).

To assess the range of behavioural thermoregulation, we conducted thermal preference experiments in a temperature Shuttlebox (Loligo Systems, Viborg, Denmark). Experimental pools were set with a static gradient of 10 °C in the cool side and 20 °C in the warm side, connected with a water bridge at 15 °C (acclimation temperature). Temperature was continually monitored and recorded by immersed temperature probes, connected to a computer-driven temperature controller and data acquisition system (DAQ-M, Loligo Systems, Viborg, Denmark). Movement in the Shuttlebox was tracked by an infrared-sensitive uEye USB 2.0 camera (IDS Imaging Development Systems GmbH, Obersulm, Germany) which connected to Shuttlesoft (v2, Loligo Systems, Viborg, Denmark) and allowed monitoring of fish movement. Following recovery from CTmax experiments (at least three days), individuals were acclimated in the Shuttlebox for 15 minutes, with entry at ‘cool’ or ‘warm’ pool randomized, to allow for exploration of the gradient. After acclimation to the experimental apparatus, the movement of each individual was tracked in 1 second intervals for 30 minutes, recording the preferred ambient water temperature (thermal preference). Individuals were assessed individually in the Shuttlebox apparatus to avoid any confounding social behaviours (*e.g.,* as seen in shoaling behaviours, Cooper et al., 2018). At the time of data collection for thermal trait experiments, all individuals were adults.

Relationships between all thermal traits separated by population and generation were assessed using the Pearson product-moment correlation with the *corrplot* (v0.84, Wei et al., 2017) and *Hmisc* (v3.14-0, Harrell, 2014) packages in R (R Core Team, 2021).

### Isolation and characterization of single nucleotide polymorphisms (SNPs)

Genomic DNA was extracted from caudal fin tissue using a phenol-chloroform-based protocol. We digested tissues overnight in digestion buffer and proteinase K at 55 °C, then performed multiple phenol-chloroform and ethanol washes to isolate the DNA. We assessed the quantity of the extracted DNA using the Quant-iT PicoGreen dsDNA assay kit (ThermoFisher Scientific, Waltham, MA, USA) and Synergy HT plate reader with the Gen5 associated software (BioTek, Winooski, VT, USA). We prepared restriction site-associated DNA (RAD) libraries (Peterson et al., 2012) using *MluCl* and *NlaIII* restriction enzymes (New England Biolabs, Ipswich, MA, USA), ligation of individual barcodes, and pooling of 48 individuals per library at equimolar concentrations. We performed a final PCR to amplify DNA and add library-specific indices to allow for pooling of multiple libraries. We sequenced three libraries at the McGill University and Génome Québec Innovation Center on one lane of Illumina HiSeq 4000 (Illumina Inc., San Diego, CA, USA).

### Assembly of the genetic linkage maps

After barcode demultiplexing and filtering out low quality reads in STACKS (Catchen et al., 2013), we removed PCR duplicates from the raw sequences and aligned these cleaned sequences to the *G. aculeatus* reference genome (Jones, Grabherr, et al., 2012) using the Burrows-Wheeler transform (Li & Durbin, 2010). Individual libraries were concatenated and filtered (Puritz et al., 2014) using *vcftools* (v3.0, Danecek et al., 2011) and split into chromosome-specific variant call format (VCF) files using *SnpSift* (Cingolani et al., 2021) to assemble the linkage maps chromosome by chromosome. We assigned markers to a linkage group with an initial logarithm of the odds (LOD) score of 3 after filtering out markers that showed high levels of segregation distortion and missing observations (> 20% missing data) in Lep-MAP3 (Rastas, 2017). Unassigned markers were subsequently added to the existing linkage group at a LOD score of 3 and a minimum size limit of 5 markers per linkage group. We ordered the markers using a minimum posterior value of 0.001 and collapsed multiple markers when the probability difference between markers was < 0.01 (Rastas, 2017). The final linkage map was phased in Lep-MAP3 (Rastas, 2017) and subset for use in R (R Core Team, 2021) with to generate a list of informative SNPs to use in subsequent analyses with the *qtl* (v1.44-9, Broman et al., 2003) package. The final linkage maps were similar across families in length and spacing between markers, though the H2_F2 map had a higher density of markers (Table S1).

### Quantitative trait loci (QTL) mapping

We analysed the four F2 families separately with the same methodology to assess the presence of QTL associated with the thermal traits and avoid the confounding effects of unique meiotic events. We calculated conditional genotype probabilities using a hidden Markov model, allowing for possible genotyping errors at a level of 0.0001 using a Kosambi mapping function with a fixed step width, prior to running genome scans with a single QTL model (Arends et al., 2014; Broman & Sen, 2009). We determined the LOD score significance thresholds for each trait through permutation tests for each family (5,000 permutations per chromosome). We pulled significant QTL above the genome-wide significance threshold (α = 0.05, Greenwood et al., 2011), calculated confidence intervals of QTL location based on nearby markers, and estimated the percent variance explained by each QTL peak marker (*PVE = 1 – 10^((−2*LOD) /n*). We identified two QTL associated with CTmin, two QTL associated with CTmax, and three QTL associated with thermal preference (Table 1, Fig. 2).

**Figure 2.**
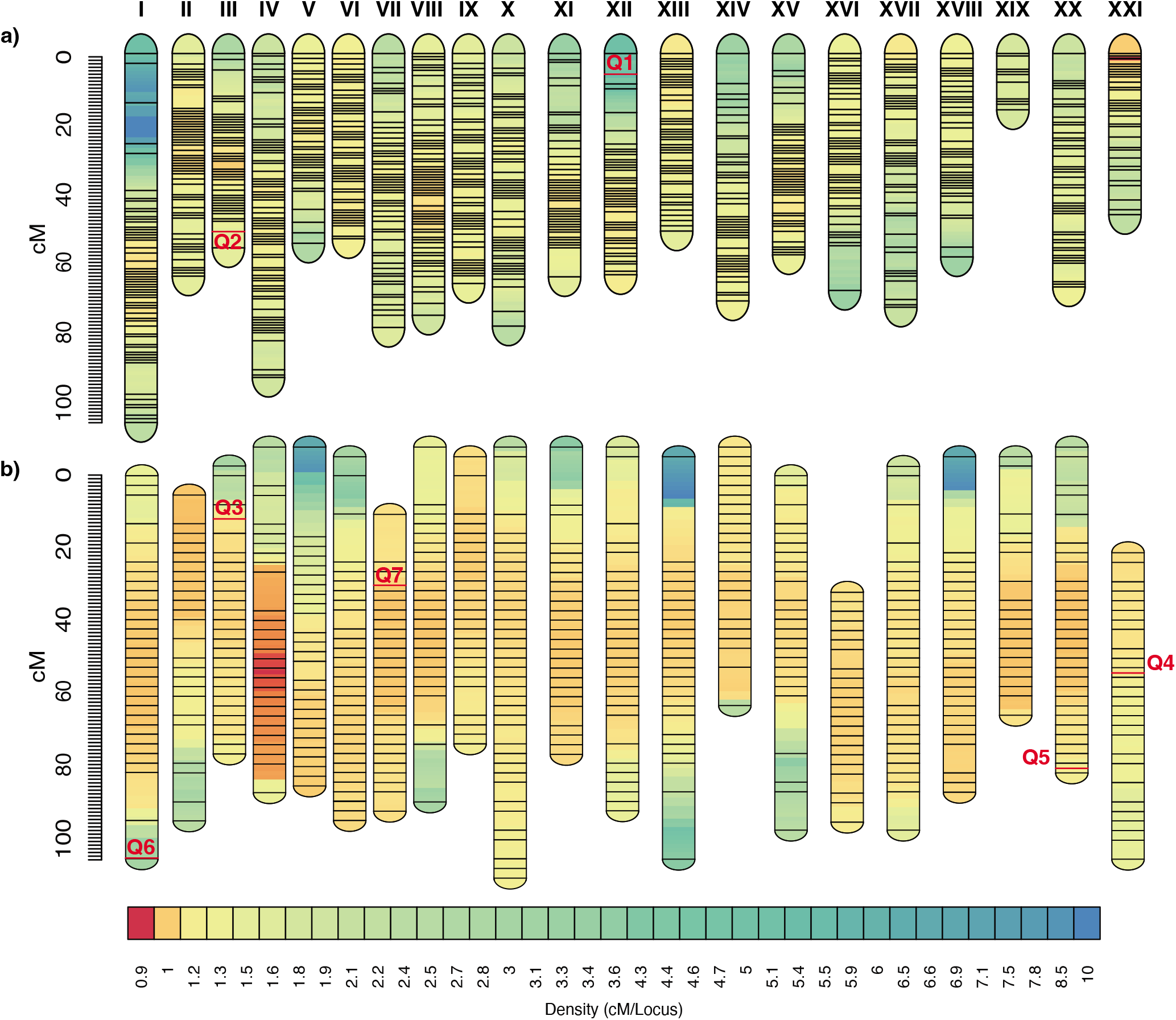
Linkage maps constructed for quantitative trait loci (QTL) analyses for a) H1 F2 hybrids (M1 x FW1) and b) H2 F2 hybrids (M1 x FW2) families, rendered with LinkageMapView (Ouellette et al., 2018), with locations of QTL indicated (Table 1).

### Environmental variables used to construct species distribution models (SDMs)

We compiled environmental data widely used in the construction of SDMs to estimate suitable habitat in both present day and end-of-century forecasts (Wiens et al., 2009), including bathymetry, sea ice extent and concentration, salinity, and sea surface temperature. We used 2014 data as our baseline year to match the forecasting baseline of the Fifth Assessment Report (IPCC, 2014). We assumed a suitable habitat range for this species in the Pacific Northwest to consist of coastal areas (water depth less than 200 m) where sea ice is never present *(i.e.,* no sea ice at the maximum extent). *G. aculeatus* has a broad salinity tolerance (Bayly, 2003; Divino et al., 2016) and salinity is not limiting in any part of the range (Zweng et al., 2013), therefore salinity was not included in the final present day or forecasted models. We obtained bathymetry data from the General Bathymetric Chart of the Oceans (GEBCO) of the British Oceanographic Data Centre (Weatherall et al., 2015), and maximum sea ice extent data from the Multisensory Analyzed Sea Ice Extent – Northern Hemisphere (MASIE-NH) product (Fetterer et al., 2010). We obtained maximum and minimum daily statistical mean sea surface temperatures (SST) from Reynolds et al. (2007) to create rasters of the mean upper quartile, lower quartile, absolute minimum, absolute maximum, and median temperatures for the baseline year (2014). We found the minimum, maximum, and median temperature for each 1 km^2^ grid cell across the range, then reduced the area based on the constraints of sea ice extent and bathymetry. Stickleback thermal trait data were used to set the limits of the ranges within the possible area delineated by sea ice free water of a suitable depth. Thermal trait measurements were based on our experimental findings reported in the present study.

In the end-of-century forecast for suitable habitat, we assumed bathymetry to be consistent with the modern scenario. In contrast, because the Arctic Ocean is projected to be predominantly free of sea ice in the summer by the end of the century (Johannessen et al., 2004), with significant end-of-century reductions in winter/spring sea ice concentration (reduced to a concentration of 0.1 at the Seward Peninsula, Johannessen et al., 2004), we conservatively set the maximum northern extent of the suitable habitat to the western tip of the Seward Peninsula (65°35’ N). The water temperatures were increased based on projections for large marine ecosystems of Northern Oceans from global climate models (Alexander et al., 2018; IPCC, 2014). Raster and maps were created in R (v4.1.2, R Core Team, 2021) using the packages *raster* (v3.5.15, Hijmans et al., 2020) and *rgeos* (v0.5.9, Roger et al., 2020).

### Trait inclusion in SDMs

We incorporated experimental data on the wild marine populations (Fig. 1c) to understand how inclusion of trait data may affect range projections under climate change. These trait-defined areas were overlain on the suitable habitat background to delineate projected presence based on thermal traits in both current day and IPCC-projected Representative Concentration Pathways (RCPs) 4.5 and 8.5. The trait values were kept constant from current day to end-of-century *(i.e.,* not changed) in the ‘no evolution’ projections. In contrast, in the ‘evolution’ projections, we allowed CTmin and CTmax to change based on empirically derived evolutionary rate estimates. We allowed CTmin to evolve at a rate of 0.63 *haldanes* estimated from a selection experiment previously conducted on populations belonging to the same admixed genetic cluster as the sampled populations (Barrett et al., 2011; Morris et al., 2018). At present, there are no empirical estimates of CTmax evolution for threespine stickleback. Thus, as a proxy we used a known estimate of CTmax evolution in fish, which was observed in zebrafish (0.19 *haldanes;* Morgan et al., 2020). Given that the evolutionary rates for CTmin and CTmax from Barrett et al. (2011) and Morgan et al. (2020) are quite high relative to other recorded evolutionary rates for phenotypic traits (particularly physiological traits; Sanderson et al., 2021), we also explored the influence of evolution on projected ranges when evolution occurred at more ‘typical’ rates. To do so, we allowed both CTmin and CTmax to evolve at a mean rate from a large meta-analysis of phenotypic trait evolution (0.14 *haldanes;* Sanderson et al. 2021).

Since the rates of evolution used here are estimates based on whole-organism tolerance (Barrett et al., 2011; Morgan et al., 2020), and trait evolution is constrained by the underlying genetic architecture of the trait (Barghi et al., 2020; Fisher, 1930; Orr, 1998; Ungerer & Riesebero, 2003), we next explored how selection acting on discrete, identifiable loci could impact evolutionary rates, and the subsequent effects on projected habitat range. To do so, we constructed models that scaled the evolutionary rate of CTmin or CTmax based on the observed genetic architecture from our QTL mapping *(i.e.,* we scaled by the percent of trait variation that is explained by each locus; ‘scaled evolution’ projections). However, an important restriction in the evolution of CTmin for all models incorporating trait evolution (‘evolution’ and ‘scaled evolution’ projections) was a hard boundary at 0 °C, with the assumption that population persistence in a sub-zero environment would require additional adaptations to CTmin improvement *(e.g.,* extreme adaptations observed in Antarctic notothenioid fishes, Cheng & Detrich, 2007; Detrich et al., 2000; Shin et al., 2014). Similarly, we defined a hard boundary for CTmax evolution at 42 °C (Morgan et al., 2018), with the assumption that the strength of selection for CTmax at this rate would result in decreased phenotypic variation and acclimatory ability leading to an upper thermal limit plateau (Gilchrist & Huey, 1999; Hangartner & Hoffmann, 2016; Morgan et al., 2020).

To quantify the differences in estimated suitable habitat under current day and end-of-century conditions, we compared areas for each warming scenario to the equivalent scenario under current conditions. Similarly, to compare the differences in evolutionary scenarios, the area of each end-of-century evolutionary trajectory was compared to either the contrasting RCP projection or scaled evolution projection. For these comparisons, we used North Pole Lambert azimuthal equal area projection for all maps, and georeferenced to known landmarks in ArcGIS v10.8 (Environmental Systems Research Institute, 2017) to calculate area from the maps generated in R (v.4.1.2, R Core Team, 2021; conversion ratio of 7873.42).

## Results

### Genetic basis of thermal traits

Stickleback exhibited a wide thermal tolerance range bounded by a mean CTmin of 1.89 °C (+/− 1.04 °C SD) and a mean CTmax of 30.1 °C (+/− 1.78 °C SD)(Fig. 1c, S2), with these physiological traits being highly correlated (*r* = 0.79, Fig. S1). These marine populations also tolerated a wide range of temperatures within which there was no observable stress response (5.0 – 25.0 °C). Thermal preference values were slightly above the acclimation temperature, but with more variance around the mean than is observed in the physiological traits (16.7 +/− 3.19 °C, Figs. 1c, S2). To determine if these measured thermal traits have a genetic basis and could therefore evolve in response to natural selection, we raised hybrid marine-freshwater F1 *(N* = 2) and F2 *(N* = 4; Fig. S3) families under common garden conditions and used these fish for genome-wide linkage map construction (Table S1) and quantitative trait loci (QTL) mapping. Using 41,840 high-quality single nucleotide variants generated from RAD sequencing, we ! identified two significant QTL for CTmin, two significant QTL for CTmax, and three significant QTL for thermal preference (Table 1, Fig. 2). The trait variance that each QTL explained ranged I from 28.9% (CTmin QTL on linkage group III) to 87.1% (thermal preference QTL on linkage group VII).

### Geographic range of projected suitable habitat

Based on sea ice extent and bathymetry alone, our present-day suitable habitat model suggests a marine range for this population from the southern Bering Sea to northern Washington state, and along the southeast Alaskan Panhandle (combined shaded area in Fig. 3a, approximately 654,122 km^2^). End-of-century IPCC projections resulted in a substantial increase in the overall suitable habitat area for stickleback, with an 873,059 km^2^ increase in total area (combined shaded area in Fig. 3b-e, approximately 1,527,181 km^2^) in association with a reduction in sea ice concentration at the northern end of the range.

**Figure 3.**
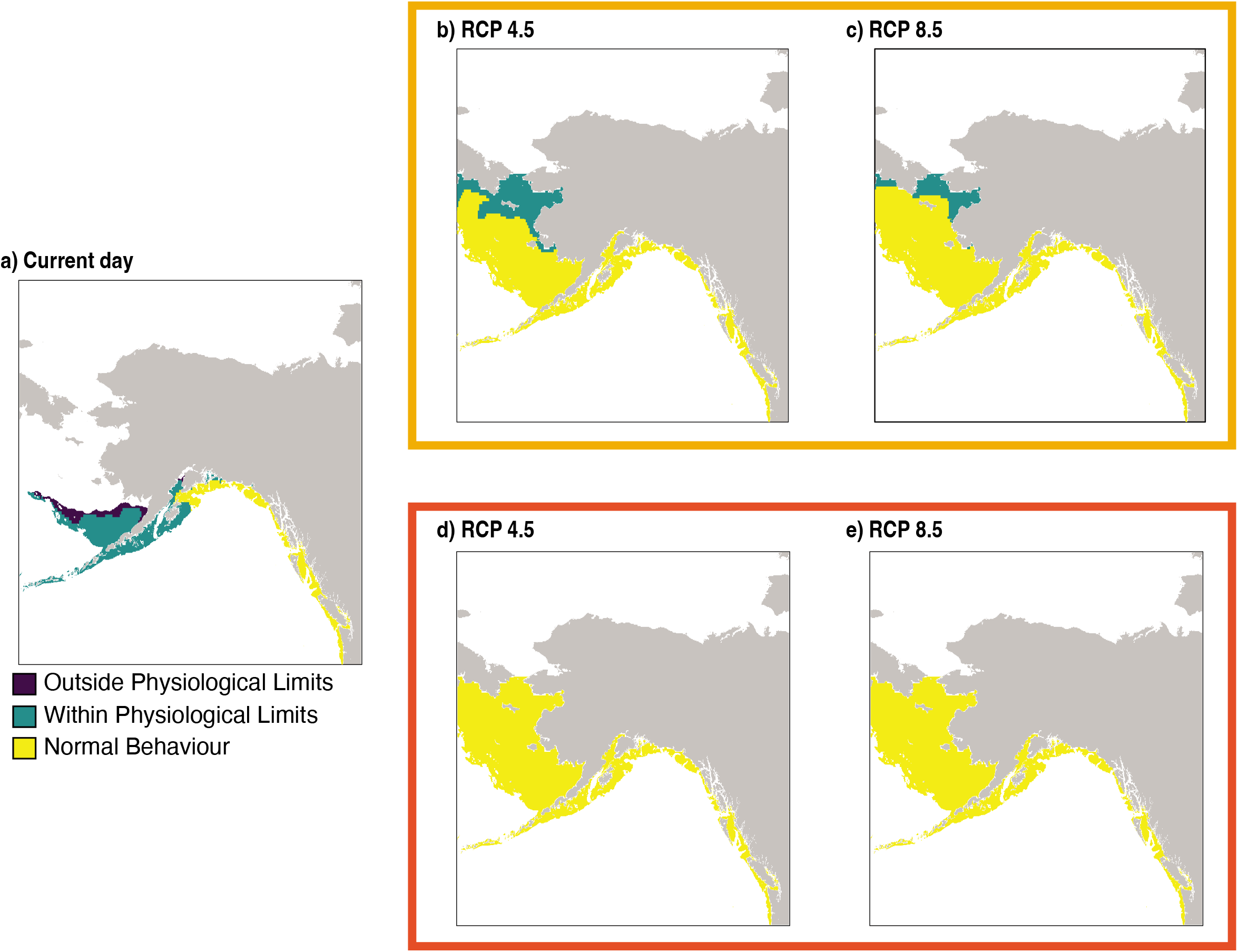
Changes in the projected range of marine threespine stickleback *(Gasterosteus aculeatus*) in the northeast Pacific Ocean as a result of incorporating thermal traits into models of a) current day environmental conditions, and under IPCC end-of-century projections RCP 4.5 and 8.5, either without trait evolution (b & c; ‘no evolution’ model, orange box) and with trait evolution at a rate of 0.63 *haldanes* for critical thermal minimum (CTmin) and a rate of 0.19 *haldanes* for critical thermal maximum (CTmax) (d & e; ‘evolution’ model, red box). Sampling locations are in the southeast area of the range presented here.

### ‘No evolution’ SDM

We used our observed values of CTmin, CTmax, and thermal preference to first inform the boundaries of three distinct environmental regions in species distribution models (SDMs): i) a ‘Normal Behaviour’ (NB) area that included environmental temperatures associated with an absence of an observable behavioural stress response (5.0 to 25.0 °C), ii) a ‘Within Physiological Limits’ (WPL) area that included environmental temperatures within the range of their measured physiological limits (1.8 to 30.1°C), and iii) an ‘Outside Physiological Limits’ (OPL) area with environmental temperatures that fall outside the measured physiological limits (below 1.8 and above 30.1 °C). When we include thermal trait data from the wild marine populations, nearly the entire range of suitable habitat was unaffected by thermal tolerance limits, apart from a slight restriction at the northern end of the range (Fig. 3a). However, when restricted to the NB area, the range becomes confined to the west of the northern tip of Kodiak Island (Fig. 3a), a limit coinciding with the northern-most known marine population in the Pacific Northwest genetic cluster (Morris et al., 2018). When temperature increases as projected by RCP 4.5, the ‘no evolution’ model projects a 1,011,949 km^2^ increase in the NB area within this newly suitable habitat (Fig. 3b) when compared to the NB area in the current day model. Under RCP 8.5, the entirety of suitable habitat area remains within tolerable limits in the ‘no evolution’ model (Fig. 3c), with a smaller proportion of the range (10.7%) falling outside of the NB area as compared to the NB area under RCP 4.5 conditions.

### ‘Evolution’ SDMs

Incorporating the evolution of CTmin and CTmax into the SDMs (‘evolution’ models) results in a large increase in the proportion of suitable habitat that falls within the NB area. At the rate of 0.63 *haldanes* for CTmin (Barrett et al., 2011) and 0.19 *haldanes* for CTmax, the entire range of the suitable habitat range falls within the NB area (Fig. 3d, 3e). These represent a 311,974 and 155,680 km^2^ increase in the NB area relative to the ‘no evolution’ models under RCP 4.5 and RCP 8.5, respectively, and an increase in 1,271,032 km^2^ in the NB area when compared to the current day model. If we only allow one of each of these traits to evolve at a time, we find that only CTmin evolution results in a change to range distributions compared to ‘no evolution’ models (Fig. S4). Reducing the rate of evolution for these traits to the mean rate of trait evolution reported in Sanderson et al. (2021) still results in all suitable habitat falling within the NB area under both RCP 4.5 and RCP 8.5. Similarly, scaling CTmin to evolve at a rate equivalent to the response expected if selection acted solely on the detected CTmin QTL on either linkage group III (PVE = 28.9%) or XXI (PVE = 53.7%) resulted in identical range distributions as the full ‘evolution’ models. Given that CTmax evolution had no impact on these range projections, we did not construct ‘scaled evolution’ models for this trait.

## Discussion

In this study, we have produced the first ‘evolution-informed’ species distribution models that incorporate empirical data for temperature-associated traits and their evolution to understand how end-of-century projections change with the inclusion of this data. We assessed physiological (critical thermal minimum, CTmin, and maximum, CTmax), and behavioural (thermal preference) thermal traits for threespine stickleback from wild marine and freshwater populations, as well as hybrid F1 and F2 families. We provide novel characterization of quantitative trait loci underlying these traits in this species, which are likely to be important genetic targets of selection under climate change. Finally, we incorporated this trait data and three empirical estimates of evolutionary rates into mechanistic niche area species distribution models under two climate change scenarios. When trait data is included in projections but held constant *(i.e.,* ‘no evolution’ models), there is an approximately 6-fold (RCP 4.5) and 7-fold (RCP 8.5) increase in the geographic area of the NB area. The geographic ranges projected for the end-of-century increased by over 7-fold when CTmin was allowed to evolve at observed trait-specific rates in the ‘evolution’ models. Additionally, when a slower rate of CTmin was used or the rate was scaled by the percent of the trait variance that is explained by individual loci (reflective of a scenario where selection acted solely on those loci), this substantial increase in NB area was still attained by the end of the century under either RCP scenario. These changes to the projected species ranges under climate change underscore the importance of incorporating behavioural as well as physiological data into SDMs (Sunday et al., 2012), as well as the key role that thermal trait evolution could play in range shifts (Buckley et al., 2010; Evans et al., 2015; Lyon et al., 2019).

Our models project a northward range expansion under climate change. This is unsurprising because climate change opens newly available thermal niche space in waters north of the current day geographic range (Alexander et al., 2018) (as is seen in the ‘no evolution’ model northward expansion, Fig. 3b & c). Northward range expansion with climate change due to increasing habitat availability has also been documented in birds (Melles et al., 2011; Rushing et al., 2020; Tingley et al., 2009; Tombre et al., 2019), plants (D’Andrea et al., 2009), other fishes (Fossheim et al., 2015; Spies et al., 2020; Yapıcı et al., 2016), and pest species (such as ticks and mountain pine beetle, Clow et al., 2017; Kurz et al., 2008; Ogden et al., 2006; Sagurova et al., 2019; Sambaraju et al., 2019), as well as in large scale analyses of diverse taxa assessing the ‘fingerprints’ of climate change impacts (Parmesan & Yohe, 2003; Platts et al., 2019). What is perhaps less expected is that our models reveal that evolution of cold tolerance, but not heat tolerance, had a substantial impact on predicted ranges under climate change, despite end-of-century climate change scenarios projecting an overall warmer, not cooler world. This result occurs because even with climate change, waters within the marine range of this stickleback population are not projected to reach temperatures that would surpass their current CTmax limit. However, there is reason to think that the relevance of cold tolerance in climate change scenarios is not specific to this system and could be generally important to consider. Climate change is leading to an increase in the frequency of extreme temperature events (IPCC, 2014; Stott, 2016), including both extreme heat and extreme cold (Herring et al., 2015, 2018, 2020), which could drive selection on both CTmin and CTmax (Buckley & Huey, 2016; Denny & Dowd, 2012; Hoffman & Sgrò, 2011; Kingsolver et al., 2011). Moreover, although northern waters are warming on average, there has been considerable spatial heterogeneity in temperature change (Walther et al., 2002). Importantly, northern waters are not predicted to be uniformly warmer under climate change (Alexander et al., 2018; IPCC, 2014) relative to the southern waters in this range, especially when seasonal variation is considered (Alexander et al., 2018). Seasonal variation and more extreme thermal events under climate change scenarios may force organisms to respond independently to both extreme heat and extreme cold in separate events (Herring et al., 2015, 2018, 2020; Walther et al., 2002), and coupled with the correlation between CTmin and CTmax (as observed here, Fig. S1), population persistence will rely on adaptation in both upper and lower critical limits. Further investigations to test the empirical rate of evolution of thermal behaviour, physiology, and the molecular underpinnings of these key traits would be well served by assessing additional subpopulations along the latitudinal gradient inhabited by stickleback to gain a more detailed understanding of these temperature-associated traits over a wider environmental range.

In interpreting the projections we present here, it is important to note that using a different methodology to assess thermal limits could result in different thresholds in the tolerance and behaviour areas (Moyano et al., 2017). In addition, in the absence of an empirical estimate of the rate of evolutionary change for CTmax in threespine stickleback, we are using a rate observed for this trait in zebrafish as a proxy, which will necessarily be inaccurate for stickleback. It is unclear whether the true rate of CTmax evolution in stickleback might be slower or more rapid than the zebrafish rate. For instance, it is possible that in threespine stickleback CTmax could evolve at similar rates as observed for CTmin, particularly given similarities in genetic architecture and the correlation between these traits in this study. Although incorporating evolution of CTmax had no impact on our projected ranges because temperatures under RCP 4.5 and RCP 8.5 never exceeded current CTmax values, we suspect this trait will be of strong importance in many other systems where populations reside in environments closer to their current upper thermal limits *(e.g.,* Deutsch et al., 2008; Morgan et al., 2019; Nati et al., 2021). Ultimately, given the dramatic temperature changes that are projected for marine systems by the end-of-century (IPCC, 2014, 2018), there is a critical need for empirical measures of evolutionary rates in thermal traits for predicting future species ranges.

While this set of mechanistic niche models builds upon current SDM work by incorporating functional traits critical to population persistence as well as varied evolutionary scenarios based on empirical measurements, there are limitations to these projections. We present niche models of geographic species ranges, but future work in this area would benefit from including a robust metric of species distribution and population genetic structure within these mechanistic geographic ranges. Deriving distributional maps from statistical algorithms that rely on species occurrence and bioclimatic variables would move beyond binary outcomes of habitat suitability and provide a map of suitable habitat that is scored as an index and could underlie the static and evolving trait areas. Species distributions that build upon these evolution-informed species ranges could be an important tool for projecting population persistence under climate change.

The efficiency of translating the selection acting on a trait into evolutionary response across generations will depend on the genetic architecture of the trait and the extent of environmental change (Dittmar et al., 2016; Orr, 2005; Rogers et al., 2012). Among-trait genetic correlations can also either speed or slow adaptive evolution depending on whether correlations are reinforcing or antagonistic to the direction of selection (Etterson & Shaw, 2001). In this case, we observe a positive correlation between CTmin and CTmax that could facilitate selection for increased temperature tolerance at both upper and lower limits. However, the empirical rates of trait evolution that we incorporated into our SDMs were estimated using change in phenotypic variation across generations (Barrett et al., 2011; Morgan et al., 2020), rather than evolution at underlying loci. As such, it is likely that some proportion of the observed phenotypic change was due to plastic responses. In our ‘scaled evolution’ models, we took a conservative approach by restricting phenotypic evolution to only occur through heritable change via the loci shown here to be associated with the trait. The large effect loci that we identified are consistent with expectations from theory that suggest prolonged bouts of adaptation with gene flow (as expected in this system, Jones, Chan, et al., 2012; Rogers et al., 2012; Schluter et al., 2010; Schluter et al., 2021; Schluter & Conte, 2009) will favour architectures characterized by fewer, larger effect, more tightly linked alleles (Griswold & Whitlock, 2003; Via et al., 2012; Yeaman & Otto, 2011; Yeaman & Whitlock, 2011). However, the effects of the QTL identified from these populations are likely overestimated, while other loci may have gone undetected (*sensu* the Beavis Effect, Beavis, 1994). These loci may have been detected in some F2 families and not in others because each mapping family has their own specific meiotic events that enable the identification of certain loci. Additionally, there are many key functional traits that are relevant to population persistence under a changing climate, and it is possible that the traits focused on here, or unexplored traits related to thermal performance, could display antagonistic pleiotropy.

Collectively, the inclusion of thermal traits and their evolution alters the projected ranges of marine threespine stickleback, with a substantial increase in the projected area that the species may occupy under climate change forecasts. Many traits are evolving in response to climate change (Eliason et al., 2011; Gómez-Ruiz & Lacher, 2019; Horton et al., 2020; Hovel et al., 2016) and SDMs that do not take trait data (and trait evolution) into account could provide less accurate predictions about future species distributions under climate change (Bush et al., 2016; Guisan et al., 2013; Huey et al., 2012; Kearney & Porter, 2004) - an issue of particular concern for species at risk and pest species undergoing range expansion (Bebber, 2015; Cullingham et al., 2011; McLeod et al., 2012). Our results provide a framework for addressing this problem, which will have critical implications for the application of these models in policy, sustainable resource management, and the protection of biodiversity in a changing climate.

## Acknowledgments

The authors acknowledge that the species collections took place on Huu-ay-aht and Sechelt First Nations traditional territories and are grateful for the opportunity to conduct their research in protected and sacred areas. We would like to thank the Bamfield Marine Sciences Centre for the resources required to conduct this research, Sam Owens for the photo in Figure 1a, Peter Peller for creating Figure 1b, Carolyn Tepolt for providing the high-resolution sea surface temperature dataset from Reynolds et al. (2007) during a government shutdown, as well as Kiyoko Gotanda for calculating and providing mean *haldane* values from the Sanderson et al. (2021) database. We would like to thank Daniel Wuitchik, Sam Yeaman, Jennifer Sunday, and Patrick Nosil for feedback on the manuscript, and the anonymous peer reviewers for improving this work. This research was funded by Discovery Grants from the Natural Sciences and Engineering Research Council of Canada to RDHB and SMR, a Canada Research Chair to RDHB, and a New Faculty Award from Alberta Innovates Technology Futures to SMR.

## Author Contributions

This study was designed by SJSW, RDHB, and SMR; fish husbandry and breeding by SJSW and TNB; experimental data collection by SJSW; DNA sequencing and initial processing by AP; bioinformatic and QTL analyses by SJSW; species distribution modelling by SJSW and SM; the manuscript was written by SJSW, RDHB, and SMR, with input from authors; the study was funded by HAJ, RDHB, and SMR.

## Data Accessibility

The data associated with this work is available as follows: ddRAD DNA sequences: available on NCBI (BioProject PRJNA795470)

Experimental data associated with thermal trait data: available on Dryad (https://doi.org/10.5061/dryad.70rxwdbz6)

Annotated code and example files: available on GitHub (https://github.com/sjswuitchik/gasAcu_qtl_sdm)

---

**Table 1.** Significant quantitative trait loci (QTL) for critical thermal minimum (CTmin), critical thermal maximum (CTmax), and thermal preference (Preference) traits with the percent of trait variance explained (PVE) for each QTL, the logarithm of odds (LOD) threshold for significance, and the LOD peak observed for each QTL.

**Figure S1.** Correlations between critical thermal minimum (CTmin), critical thermal maximum (CTmax), and thermal preference for threespine stickleback (*Gasterosteus aculeatus*).

**Figure S2.** Thermal preference (green), critical thermal minimum (CTmin, blue) and maximum (CTmax, red) measurements for threespine stickleback (*Gasterosteus aculeatus*) from wild freshwater (FW) populations, pure F1 freshwater (FW*F1), wild marine (M), and pure F1 marine (M*F1) families raised in a common garden under a constant thermal environment.

**Figure S3.** Thermal preference (green), critical thermal minimum (CTmin, blue) and maximum (CTmax, red) measurements for threespine stickleback (*Gasterosteus aculeatus*) from hybrid marine-freshwater F2 families raised in a common garden under a constant thermal environment.

**Figure S4.** Changes in the projected range of marine threespine stickleback *(Gasterosteus aculeatus*) in the northeast Pacific Ocean compared to a) current day range as a result of incorporating the evolution of only critical thermal minimum (CTmin, b & c; orange box) at a rate of 0.63 *haldanes* and only critical thermal maximum (CTmax, d & e; red box) at a rate of 0.19 *haldanes* under IPCC end-of-century projections RCP 4.5 and RCP 8.5

**Table S1.** Summaries of linkage maps constructed for quantitative trait loci (QTL) analyses.

