## Supplementary figures and images for "Evolution of thermal physiology alters the projected range of threespine stickleback under climate change"

### Supplemental Figure 1

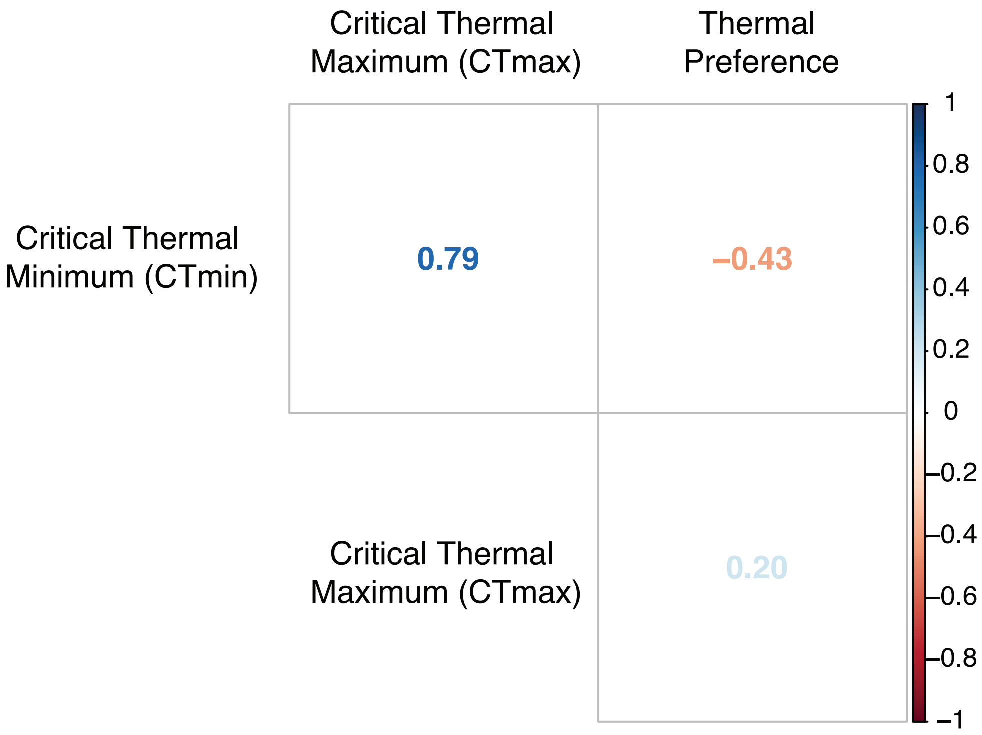

### Supplemental Figure 2

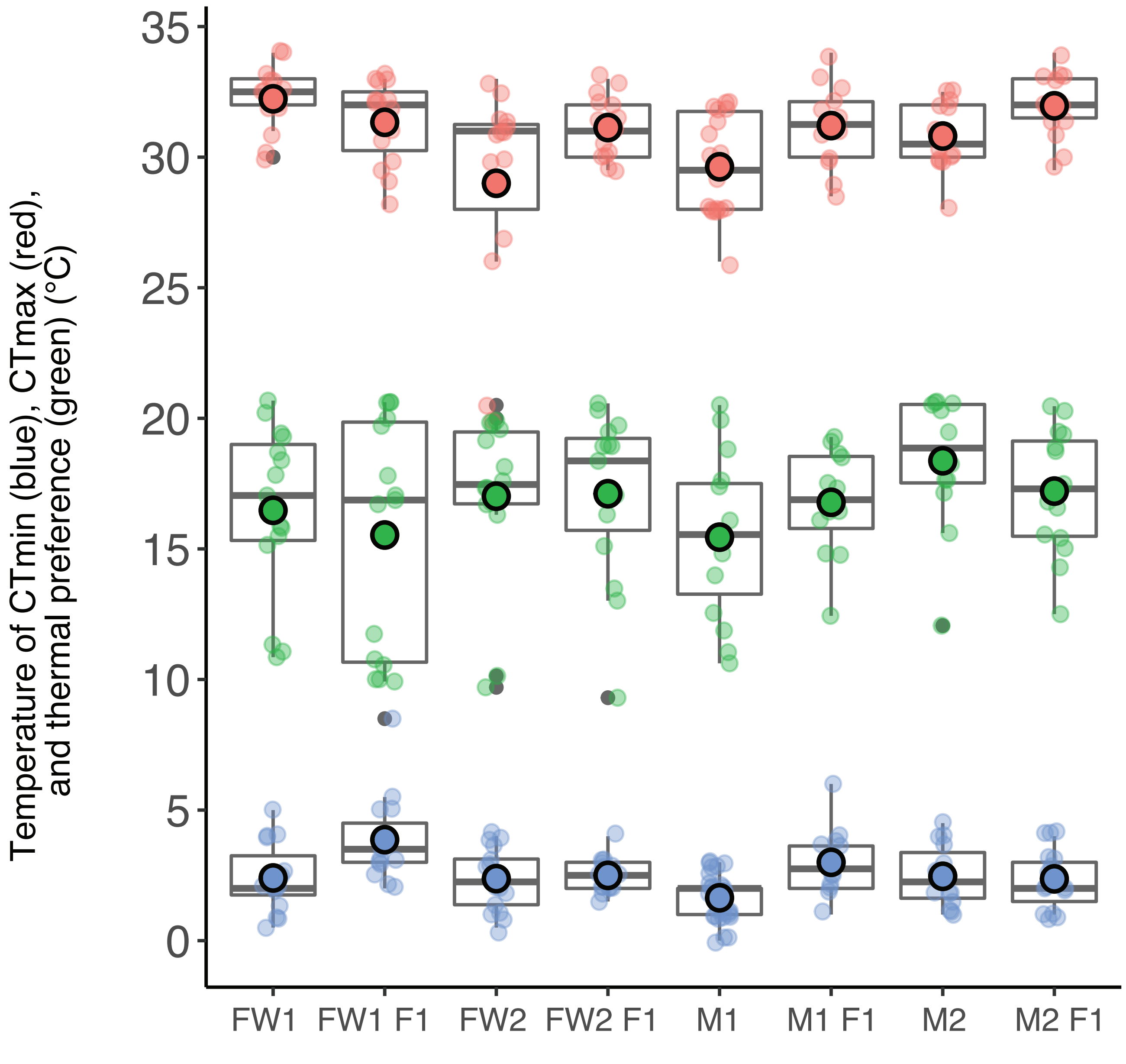

### Supplemental Figure 3

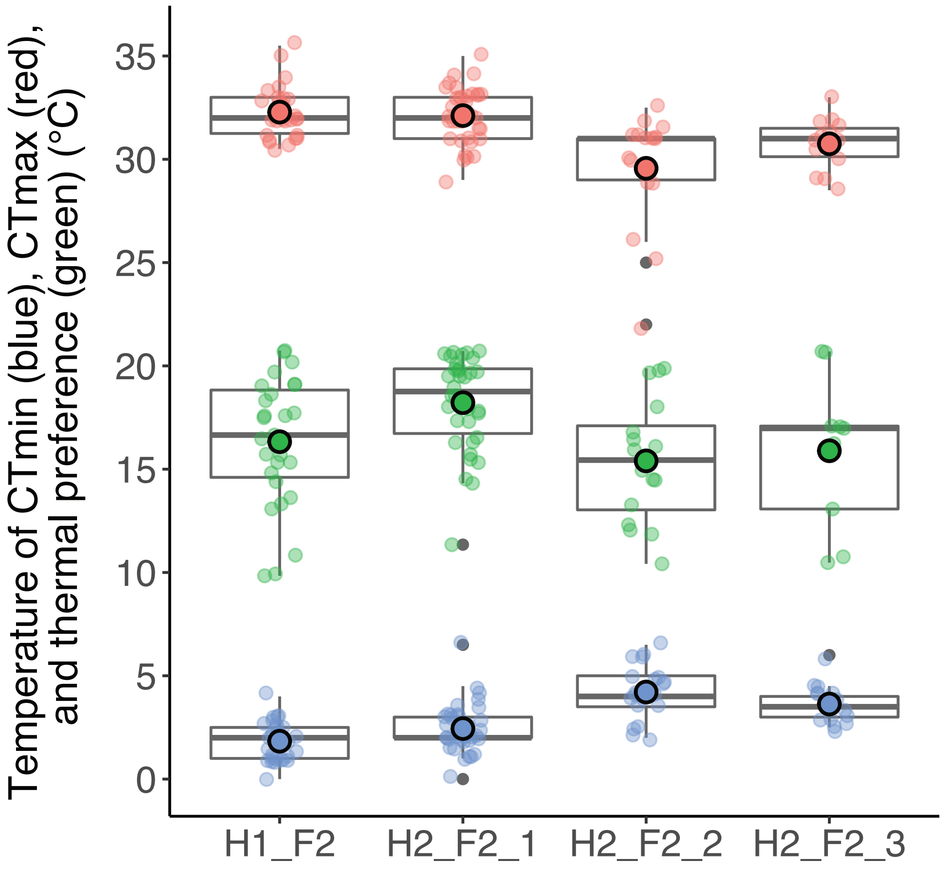

### Supplemental Figure 4

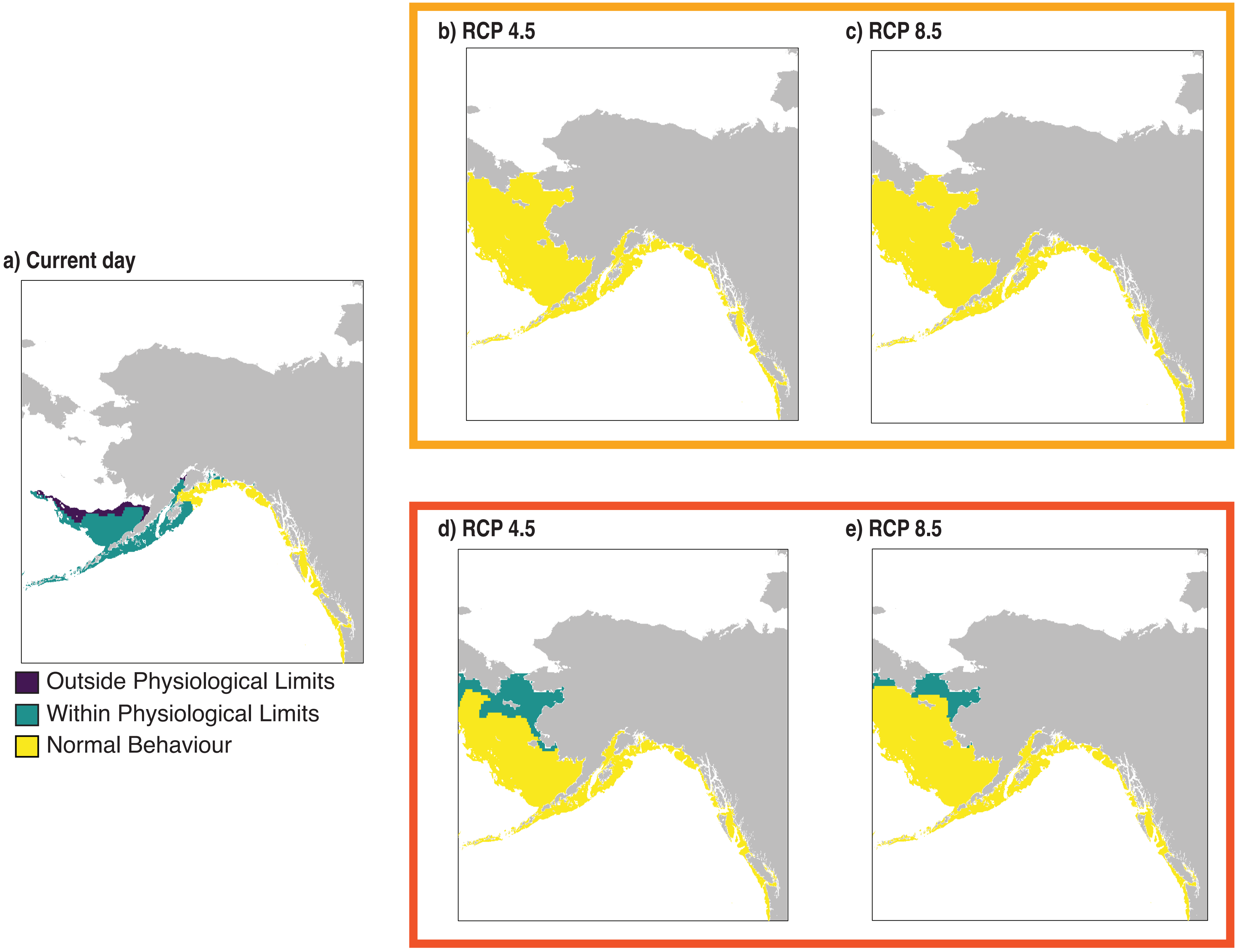
