## Supplementary material for "Evolution of thermal physiology alters the projected range of threespine stickleback under climate change": Table 1

| QTL Name | Family | Phenotype | Linkage Group | Position (cM) | 95 % CI | PVE (%) | LOD threshold (5%) | LOD peak |
| --- | --- | --- | --- | --- | --- | --- | --- | --- |
| Q1 | H1_F2 | CTmax | XII | 4.56 | 0.000 – 11.38 | 45.2 | 2.71 | 2.87 |
| Q2 | H1_F2 | Preference | III | 50.04 | 43.21 – 70.80 | 53.3 | 2.78 | 3.64 |
| Q3 | H2_F2_1 | CTmin | III | 11.57 | 6.347 – 52.62 | 28.9 | 2.55 | 2.63 |
| Q4 | H2_F2_2 | CTmin | XXI | 55.67 | 37.01 – 66.87 | 53.7 | 2.75 | 3.01 |
| Q5 | H2_F2_2 | CTmax | XX | 81.36 | 73.12 – 88.48 | 83.3 | 5.57 | 7 |
| Q6 | H2_F2_2 | Preference | I | 108.2 | 83.54 – 109.2 | 69.3 | 3.27 | 4.62 |
| Q7 | H2_F2_3 | Preference | VII | 31.48 | 24.02 – 53.87 | 87.1 | 3.61 | 5.78 |
