## Supplemental Table 1 for "Evolution of thermal physiology alters the projected range of threespine stickleback under climate change"

**Table S1.** Summaries of linkage maps constructed for quantitative trait loci (QTL) analyses.

| Map | Nmarkers | Length (cM) | Average Maximum Spacing |
| --- | --- | --- | --- |
| H1_F2 | 6571 | 1678.7 | 8.98 |
| H2_F2 | 9930 | 1995.3 | 9.36 |
